# The net charge of the K-loop regulates KIF1A superprocessivity by enhancing microtubule affinity in the one-head-bound state

**DOI:** 10.1101/2022.08.21.504701

**Authors:** Taylor M. Zaniewski, William O. Hancock

**Affiliations:** Departments of Chemistry and Biomedical Engineering, Pennsylvania State University, University Park, Pennsylvania, USA

## Abstract

KIF1A is an essential neuronal transport motor protein in the kinesin-3 family, known for its superprocessive motility. We determined that superprocessivity of KIF1A dimers originates from a unique structural domain, the lysine rich insertion in loop-12 termed the ‘K-Loop’, which enhances electrostatic interactions between the motor and the microtubule. In 80 mM PIPES buffer, replacing the native loop-12 of KIF1A with that of kinesin-1, resulted in a 6-fold decrease in run length, and adding additional positive charge to loop-12 enhanced the run length. Interestingly, swapping the KIF1A loop-12 into kinesin-1 did not enhance its run length, consistent with the two motor families using different mechanochemical tuning to achieve persistent transport. To investigate the mechanism by which the KIF1A K-loop enhances processivity, we used microtubule pelleting and single-molecule dwell times assays in ATP and ADP. First, the microtubule affinity was similar in ATP and in ADP, consistent with the motor spending the majority of its cycle in a weakly-bound state. Second, the microtubule affinity and single-molecule dwell time in ADP were 6-fold lower in the loop-swap mutant compared to wild type. Thus, the positive charge in loop-12 of KIF1A enhances the run length by stabilizing the motor binding in its vulnerable one-head-bound state. Finally, through a series of mutants with varying positive charge in the K-loop, we found that the KIF1A processivity is linearly dependent on the charge of loop-12.

## Introduction

KIF1A is a fast and superprocessive neuronal transport motor protein in the kinesin-3 family that is responsible for the delivery of synaptic vesicle precursors from the soma to the distal end of the axon, among other tasks.^1–5^ Genetic mutations lead to KIF1A Associated Neurological Disorders (KAND), rare and often misdiagnosed afflictions.^5–14^ The category of KAND is vast, with 113 variants that have been identified in humans so far, making therapeutic attempts difficult.^14^ Therefore, approaching treatments more systematically requires a deeper understanding of the fundamental mechanochemical mechanism of KIF1A.

A recent biochemical dissection of the KIF1A chemomechanical cycle concluded that: i) both ATP binding and hydrolysis are required to trigger tethered-head binding, ii) rear head detachment is fast and may contribute to the fast stepping rate, and iii) the motor spends the majority of its stepping cycle in a post-hydrolysis, one-head-bound (1HB) state.^15^ This ability to remain associated with the microtubule in a “weakly-bound” state enhances the motor processivity by ensuring that the motor remains bound to the microtubule sufficiently long for the tethered head to complete its step and bind to the next site.^16–18^ This property of remaining in a vulnerable one-head-bound state also provides an explanation for the propensity of KIF1A to detach readily under load.^19–22^ A recent optical trapping study that used a three-bead geometry to minimize forces oriented perpendicular to the microtubule concluded that under load KIF1A is able to sustain loads by rapidly rebinding to the microtubule following disengagement.^20^ Because KIF1A does not achieve high microtubule affinity by maximizing the duration spent in a strong binding two-head-bound state, we proposed that the source of its processivity is a combination of a relatively slow detachment rate from the post-hydrolysis 1HB state and faster tethered-head attachment rate compared to kinesin-1 and −2.^15^ The goal of the current study was to define the structural elements of KIF1A that underlie this kinetic tuning.

The kinesin-3 family is known for a unique structural domain called the K-Loop. This domain is a stretch of Loop-12 in which KIF1A contains six lysine residues and kinesin-1 and −2 only have one lysine.^23^ In early work it was determined that the electrostatic interaction of the K-Loop with the microtubule facilitates diffusive motion, leading to monomeric motility of KIF1A.^23–25^ The enhanced positive charge of KIF1A in this region is thought to interact with the glutamate rich C-terminal tail of tubulin, termed the ‘E-hook’.^26,27^ However, because these domains are disordered, and their structures and interactions not resolved by X-ray crystallography or CryoEM,^26,28,29^ biochemical and single-molecule investigations are vital for understanding this interaction.

In the literature, there are conflicting reports regarding the role of the K-loop in KIFA superprocessivity, with some studies indicating that the K-loop is responsible for KIF1A superprocessivity,^27,30^ and others suggesting that the K-loop plays no role in the processivity of KIF1A and only increases its microtubule on-rate.^31^ This lack of consensus in the field has led to considerable confusion regarding the role of the K-loop in long distance intracellular transport. In the present study, we used single-molecule microscopy and stopped-flow biochemical experiments to clarify the role of loop-12 in KIF1A motility. By using a series of mutations, buffers, and experimental approaches, we determined that at near physiological ionic strength the K-loop enhances the processivity of KIF1A by strengthening the microtubule affinity of a vulnerable one-head-bound state in the KIF1A chemomechanical cycle. Additionally, this functionality cannot be transferred to kinesin-1, highlighting how different motor families use distinct biochemical tuning to achieve fast and processive motility.

## Results

### Influence of the neck-coil and coiled-coil domains on KIF1A motility

To understand how different structural elements of KIF1A contribute to its motility and to reconcile conflicting results in the literature, we designed constitutively active constructs of KIF1A that included different distal coiled-coil domains to ensure stable KIF1A dimers and included or excluded the native neck-coil domain of KIF1A. All of the KIF1A constructs are based on a truncated and constitutively active KIF1A dimer that has been used in a number of published studies.^15,27,31,32^ The ‘neck-coil’, defined as residues 369-393 of *Rattus norvegicus* KIF1A, is a short coiled-coil domain that is immediately distal to the disordered neck linker domain and is involved in dimerization of the two motor domains.^33,34^ Based on other members of the kinesin-3 family, it is thought that in full-length KIF1A, coiled-coil 1 folds back on the neck-coil and forms an antiparallel coiled-coil that separates the two heads and inhibits motor activity.^32,33^ It was shown previously that the neck-coil alone is insufficient to stably dimerize KIF1A,^32–34^ and that this could be rectified by adding a leucine zipper (LZ) downstream of the neck-coil.^19,27,32^ A second approach to dimerizing diverse kinesins has been to fuse the motor and neck linker domains to the coiled-coil domain of *D. melanogaster* kinesin-1.^15,17,35–37^

To directly compare how these different dimerization strategies affect KIF1A motility, we designed three dimeric, GFP-tagged *Rattus norvegicus* KIF1A constructs, as follows. The first construct consisted of KIF1A residues 1-393 followed by a leucine zipper domain and GFP, which we refer to as 1A-LZ (Fig. 1A). This construct matches those used in a number of published studies, albeit with different C-terminal tags.^19,27,30,32^ We made two other constructs that achieved stable dimerization through the *Drosophila melanogaster* kinesin heavy chain (KHC) neck-coil and coiled-coil 1 domains (345-560). The first construct, which includes the KIF1A neck-coil, fused KIF1A(1-393) to residues 345-560 of KHC and a C-terminal GFP, which we refer to as 1A_393_ (Fig. 1A-B). The second construct, which does not include the KIF1A neck-coil and was used in a previous study,^15^ fused KIF1A(1-368) to residues 345-560 of KHC and a C-terminal GFP. We refer to this as 1A_368_ (Fig. 1A).

**Figure 1:**
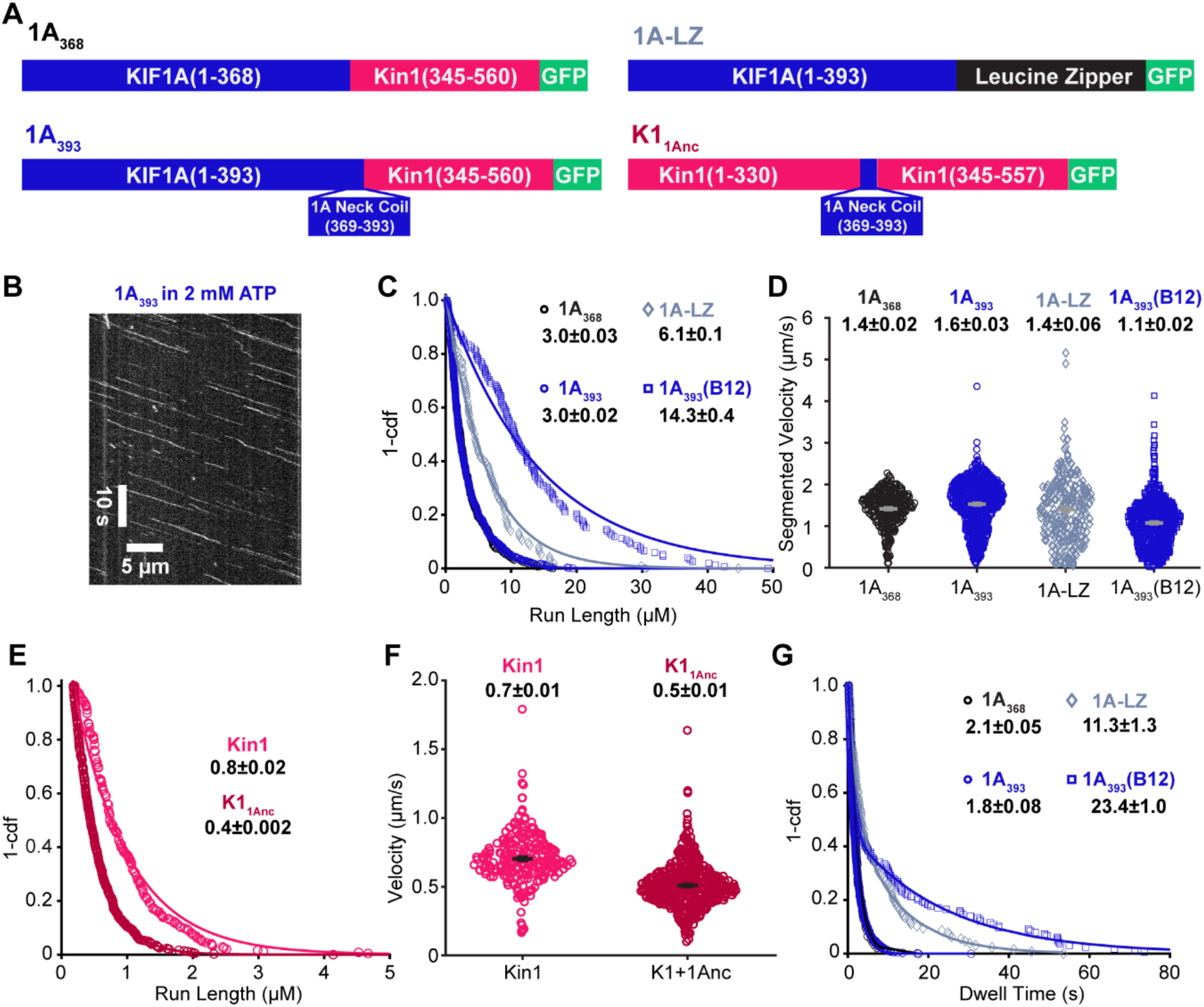
Dimerization domain and buffer ionic strength impact KIF1A motility. **A,** Diagrams of the protein constructs used in this study. KIF1A refers to *Rattus norvegicus* KIF1A; Kin1 refers to *Drosophila melanogaster* kinesin heavy chain. **B,** Example kymograph of 1A_393_ in 2 mM ATP and BRB80. Scale bars are 10 sec and 5 μm. **C,** Run length distribution of 1A_368_ (black circles), 1A_393_ (blue circles), 1A_393_-LZ (navy diamonds), and 1A_393_ in BRB12 (blue squares). Single exponential fits give run lengths of 3.0 ± 0.03, 3.0 ± 0.02, 6.1 ± 0.1, and 14.3 ± 0.4 μm, respectively (fit ± 95% confidence interval). **D,** Velocity distribution of 1A_368_ (black circles), 1A_393_ (blue circles), 1A_393_-LZ (navy diamonds), and 1A_393_ in BRB12 (blue squares). Average segmented velocities are 1.4 ± 0.02, 1.6 ± 0.03, 1.4 ± 0.06, 1.1 ± 0.02 μm/s, respectively (mean ± SEM). **E,** Run length distribution of Kin1 (pink) and K1_1Anc_ (red). Single exponential fits give run lengths of 0.8 ± 0.02, and 0.4 ± 0.002 μm, respectively (fit ± 95% confidence interval). **F,** Velocity distribution of Kin1 (pink) and K1_1Anc_ (red). Average velocities are 0.7 ± 0.2 and 0.5 ± 0.2 μm/s, respectively (mean ± SD). **G,** Dwell time distributions in 2 mM ADP. Single exponential fits to 1A_368_ (black circles) and 1A_393_ (blue circles) give dwell times of 2.1 ± 0.05 and 1.8 ± 0.08 s, respectively. 1A_393_-LZ (navy diamonds) and 1A_393_ in BRB12 (blue squares) were fit with biexponentials. 1A_393_-LZ had dwell times of 2.0 ± 0.3 and 11.3 ± 1.3 s, and 1A_393_ in BRB12 had dwell times of 1.1 ± 0.04 and 23.4 ± 1.0 s; the longer duration for each is presented in the figure.

To reconcile the behavior of disparate constructs in the literature, the first question we addressed was whether the dimerization strategy affected the KIF1A run length in 80 mM PIPES (BRB80) buffer. We found that 1A-LZ had a 2-fold longer run length than 1A_393_ (6.1 ± 0.1 and 3.0 ± 0.02 μm, respectively; Fig. 1C with raw data in Fig. S1). We propose that this difference in processivity is due to the differential charge in the coiled-coil domains rather than differences in interhead coordination between the two constructs. The net charge of coil-1 of KIF1A, which these dimerization domains replace, is −6; the net charge of the kinesin-1 neck-coil (345-405) is −3; and the net charge of the leucine zipper is neutral (Fig. S2). It was shown previously that adding positive charge to the kinesin-1 coiled-coil enhances its run length and adding negative charge diminishes its run length.^38^ Thus, we interpret the longer run length of 1A-LZ to be due to enhanced interaction between the coiled-coil domain and the negatively charged C-terminus of tubulin. One prediction of this hypothesis is that the motor off-rate in ADP, which induces a weak-binding state that doesn’t involve coordinated activities of the two heads, should be similarly affected by the electrostatic interactions between the coiled-coil and the microtubule. To probe this question, we used TIRF microscopy to measure the microtubule dwell time of the two constructs in saturating ADP. Consistent with the run length differences, 1A_393_-LZ had a ~3-fold longer dwell time than 1A_393_ (Fig. 1F). Thus, because the negative net charge of the Kin1 neck-coil sequence better matches the native KIF1A, and because this domain has been used in a body of our previous work, we focused our efforts on the 1A_393_ construct.

Because the KIF1A neck-coil domain has been shown to be insufficient to form a stable dimer on its own,^32^ it is possible that even in constructs dimerized by fusing a distal coiled-coil domain, the neck-coil does not form a stable connection between the two motor domains. This ‘breathing’ of the neck-coil could affect the interhead coordination necessary for processivity, consistent with previous work that showed extending the length of the kinesin neck linker, which connects each motor domain to their shared neck-coil, reduced motor run lengths.^34,39–41^ To test whether the KIF1A neck-coil plays a role beyond dimerization, we used TIRF microscopy in BRB80 buffer to compare the run length of 1A_393_, which includes the neck-coil, to 1A_368_, which lacks the neck-coil (Fig. 1A). The two constructs had similar run lengths, indicating that the sequence and stability of the KIF1A neck-coil has no effect on its processivity (Fig. 1B, D). Consistent with the run lengths, the microtubule dwell times of the two motors in 2 mM ADP, which probes the affinity in the weak-binding state, were also similar (Fig. 1G). We hypothesized that if there is any reversible dimerization in the KIF1A neck-coil, then it may have a stronger effect on kinesin-1, which has a shorter neck linker domain and thus a stiffer connection between the two heads than KIF1A.^17,40–43^ To test this, we used *Drosophila* kinesin-1 truncated at residue 557 and fused to GFP, which has been used in numerous previous studies, and is referred to as Kin1 here.^15,17,35,36^ The KIF1A neck-coil domain (KIF1A 369-393) was inserted just upstream of Ala345 at the start of the neck-coil of Kin1 to generate K1_1Anc_ (Fig. 1A).^44^ In single-molecule assays, the Kin1 run length was 0.8 ± 0.02 μm/s (Fig. 1E), which is consistent with previous work and is four-fold shorter than KIF1A (Fig. 1C).^17^,^35^,^36^ K1_1Anc_ had a ~2-fold shorter run length than Kin1 (0.4 ± 0.002 μm; Fig. 1D) and also had a slightly slower velocity (0.5 ± 0.2 μm/s versus 0.7 ± 0.2 μm/s; Fig. 1E). It is unlikely that the shorter run length of K1_1Anc_ results from differences in positive charge in the neck-coil domain, because both the kinesin-1 and KIF1A neck-coil domains are negatively charged (−3 for Kin1 and −2 for KIF1A; Fig. S2). Instead, the shorter run length of K1_1Anc_ is consistent with the KIF1A neck-coil dimerizing only weakly and acting as an extended neck linker in kinesin-1, loosening the connection between the two motor domains and reducing its performance.^17,40–42^ In light of this, it is surprising that replacing the KIF1A neck-coil with the more stable kinesin-1 neck-coil does not enhance the KIF1A run length. However, the native neck linker domain of KIF1A is longer than that of kinesin-1,^43^ and so one possibility is that stabilizing the KIF1A neck region by adding the kinesin-1 coiled-coil is not sufficient to establish a tight connection between the two heads. It follows that the superprocessivity of KIF1A does not result from tight mechanical connection between the heads to achieve coordinated stepping, but rather from other aspects of the motor’s chemomechanics.

### Influence of ionic strength on KIF1A motility

To better understand the role of electrostatic interactions in KIF1A motility and to reconcile diverse studies across the literature, we investigated the effect of buffer ionic strength on KIF1A motility. We chose two buffers commonly used in the literature: BRB80, which contains 80 mM PIPES and has a 173 mM ionic strength, and BRB12, which contains 12 mM PIPES and has a 26 mM ionic strength (both buffers also include 1 mM MgCl_2_, 1 mM EGTA, pH 6.9). Although much of the published work on KIF1A was performed in BRB12,^19,27,31,32^ the ionic strength of BRB80 (173 mM) is closer to the ~200 mM ionic strength estimated in cells.^45^ The low ionic strength of BRB12 is expected to maximize electrostatic interactions between motors and microtubules; for instance, the Debye length in BRB80 is 0.7 nm in BRB80 and 2 nm in BRB12.^46^

Consistent with enhanced electrostatic interactions, we found that 1A_393_ had a nearly 5-fold longer run length in BRB12 compared to BRB80 (14.3 ± 0.4 μm versus 3.0 ± 0.02; Fig. 1C). Additionally, we observed increased pausing behavior at low ionic strength, as reported previously.^27^ Thus, we quantified the segmented velocity (Fig. 1D) and found that the velocity decreased from 1.5 μm/s in BRB80 to 1.1 μm/s in BRB12. The longer run length and slower velocity correspond to a substantially higher affinity of KIF1A for microtubules in BRB12 buffer. To compare the microtubule affinity of KIF1A in its weak-binding ADP state in these different buffers, we measured the microtubule dwell time in 2 mM ADP and found that the dwell time in BRB12 was 10-fold longer than in BRB80 (Fig. 1F). Thus, the enhanced run length of KIF1A in BRB12 can be explained by the reduced charge shielding at lower ionic strength enhancing the electrostatic interaction between the microtubule and KIF1A in the weakly-bound ADP state. Because these enhanced electrostatic interactions in BRB12 may mask other aspects of KIF1A mechanochemistry, we focused our efforts on characterizing KIF1A in BRB80, which is closer to physiological ionic strength.

### K-loop regulates the run length of KIF1A but not Kin-1

Earlier studies of KIF1A monomers found that the positively charged Loop-12 of KIF1A enhances its microtubule affinity,^23^ however subsequent studies of KIF1A dimers have reported Loop-12 does not contribute to superprocessivity.^31^ To test whether the K-Loop is responsible for the superprocessivity of our KIF1A construct in BRB80 buffer, we examined the properties of a ‘swap’ construct, in which the native loop-12 of KIF1A is removed and replaced with the kinesin-1 loop-12, KIF1A(1-393)-Kin1Loop12-K560-GFP (referred to as 1A-K1L12; Fig. 2A).^23,31^ Swapping out the KIF1A Loop-12 caused a 6-fold decrease in the run length and a slight decrease in velocity, consistent with Loop-12 being the determinant of KIF1A superprocessivity (Fig. 2 B-C and Fig. S3).

**Figure 2:**
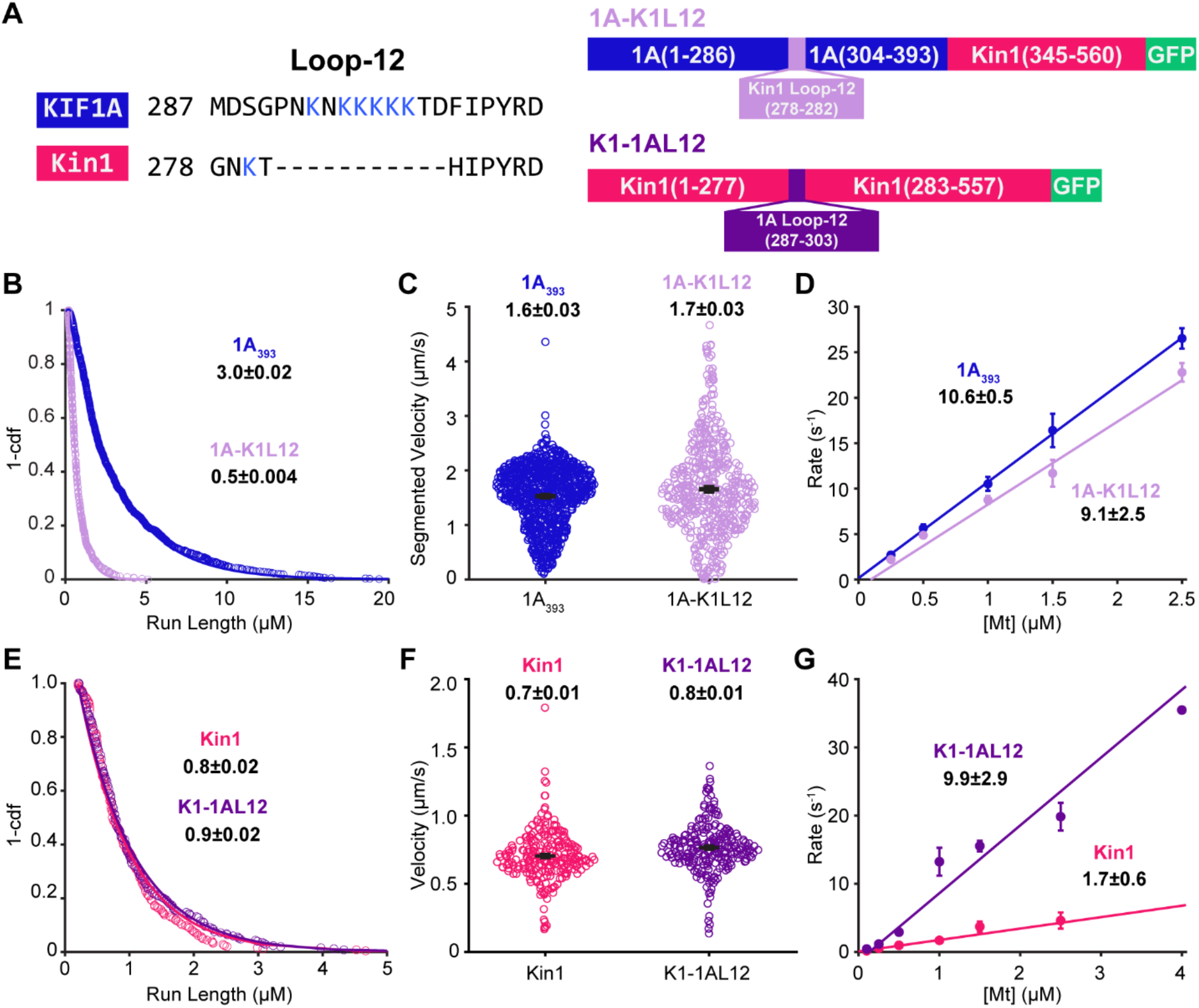
K-Loop is the source of KIF1A superprocessivity. **A,** Diagrams of the loop swap constructs. **B,** Run length distribution of 1A_393_ (blue circles) and 1A-K1L12 (light purple circles) in BRB80 and 2 mM ATP. Values are single exponential fit ± 95% confidence interval. **C,** Velocity distribution of 1A_393_ (blue circles) and 1A-K1L12 (light purple circles) in BRB80 and 2 mM ATP. Average segmented velocities values are mean ± SEM. P<0.0001 **D,** Bimolecular on-rates of 1A_393_ (blue circles) and 1A-K1L12 (light purple circles), using motors lacking coiled-coil 1 and GFP (see Methods). Linear fits give 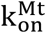 (fit ± 95% confidence interval). Points are mean ± SEM. **E,** Run length distribution of Kin1 (pink circles) and K1-1AL12 (dark purple circles) in BRB80 and 2 mM ATP. Values are single exponential fit ± 95% confidence interval. **F,** Velocity distribution of Kin1 (pink circles) and K1-1AL12 (dark purple circles) in BRB80 and 2 mM ATP. Values are mean ± SEM. P<0.0001 **G,** Bimolecular on-rates of Kin1 (pink circles) and K1-1AL12 (dark purple circles). Linear fits give 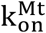 (fit ± 95% confidence interval). Points are mean ± SEM.

If Loop-12 is enhancing KIF1A processivity by enhancing electrostatic interactions with the microtubule, then in principle swapping it into kinesin-1 should enhance the run length, and previous work found this to be the case in low ionic strength buffer.^31^ To test this question in BRB80 buffer where physiological charge shielding is expected, we made a ‘reverse swap’ mutant, in which the KIF1A loop-12 was inserted into kinesin-1, replacing the native loop-12 (referred to as K1-1AL12; Fig. 2A). Surprisingly, inserting the positively charged KIF1A K-loop into Kin-1 did not impact the velocity or run length (Fig. 2E-F; see Fig. S3 for representative kymographs). Thus, due to differences in the microtubule binding interface and/or kinetic tuning of kinesin-1, adding the positively charged K-loop into kinesin-1 does not enhance its processivity in near physiological ionic strength buffer.

To further investigate how positive charge in loop-12 differentially affects KIF1A and kinesin-1, we compared the effects of the loop swap on the microtubule on-rate of the two motors. Using stopped-flow, the bimolecular on-rate 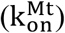 was measured by incubating the motors with fluorescently labeled mant-ADP in BRB80 and flushing the solution against varying concentrations of taxol-stabilized microtubules in 2 mM ADP. Upon mixing, the motors bind to the microtubule and release the mant-ADP, which results in a fall in fluorescence. Because at low microtubule concentrations mant-ADP release is rate limited by microtubule binding, a linear fit of the rates to the microtubule concentration yields the bimolecular on-rate.^15,42,47,48^ Notably, swapping the K-loop out of KIF1A had little effect on the bimolecular on-rate of KIF1A, contrasting with its effect on the run length (Fig. 2D). However, swapping the KIF1A loop-12 into kinesin-1 caused a 5-fold increase in the on-rate (Fig. 2G).

### Positive charge in the KIF1A K-loop enhances processivity by decreasing the rate of detachment from a vulnerable 1HB state

To investigate the mechanism by which the K-Loop enhances KIF1A processivity, we quantified key rate constants in the KIF1A chemomechanical cycle that determine motor run length. Previous work established that kinesin processivity is determined by a kinetic race as the motor takes a forward step, as follows.^17,18^ Following ATP hydrolysis, kinesin is in a vulnerable one-head-bound state that can resolve either by the tethered head completing a forward step by binding to the next tubulin binding site and transitioning to a tight binding state, or by the motor dissociating from the microtubule and terminating the run (Fig. 3A).^15,17^ Thus, in principle the K-loop could enhance KIF1A processivity by some combination of increasing the on-rate that the tethered head binds to its next binding site (k_on_^TH^) and decreasing the rate that the bound head detaches from the microtubule (k_detach_).

**Figure 3:**
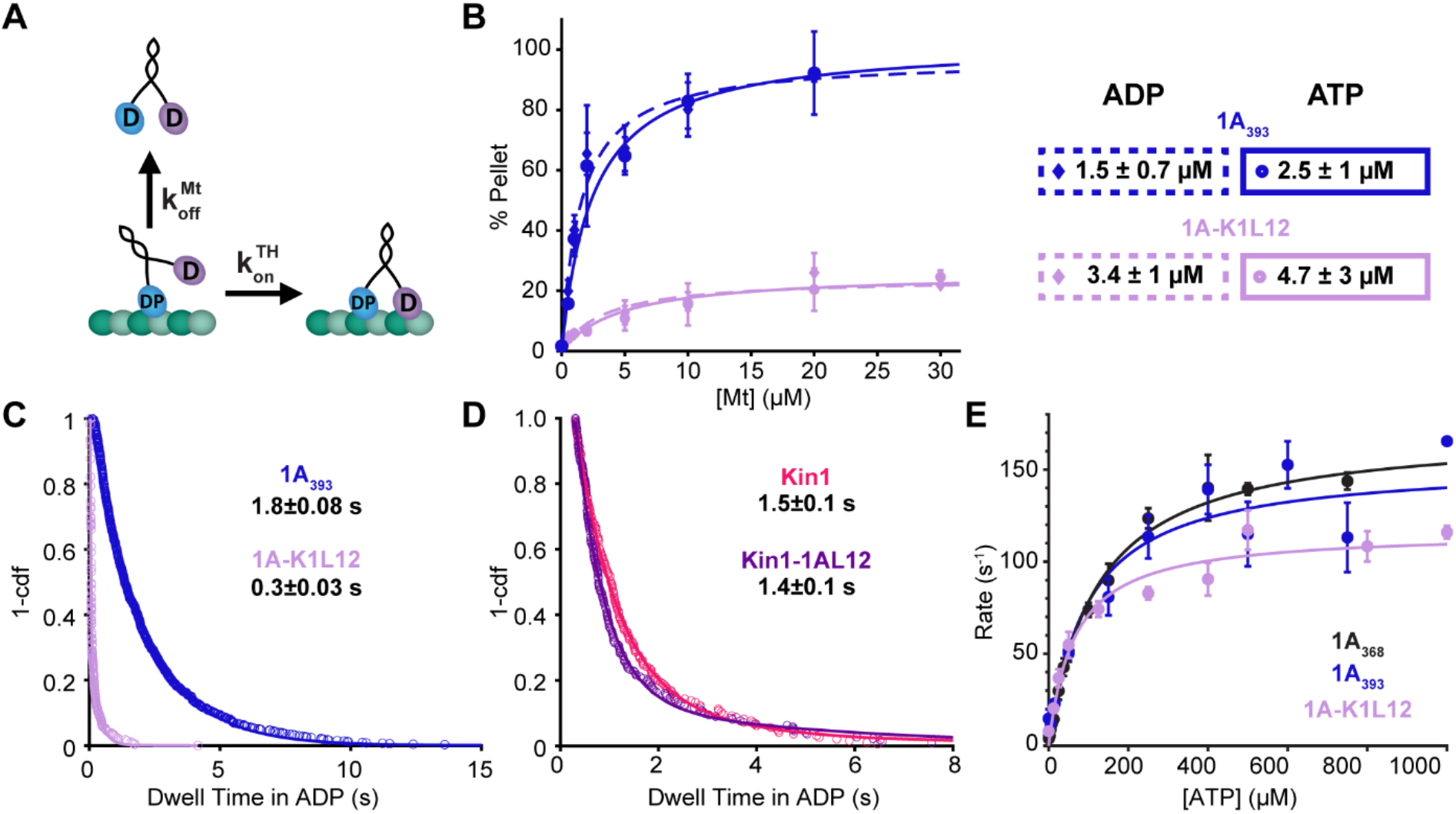
K-Loop regulates KIF1A superprocessivity via detachment rate in the kinetic race. **A,** Diagram of transitions involved in the Kinetic Race, the tethered-head attachment rate 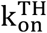 and the microtubule detachment rate k_detach_. D, ADP; DP, ADP+Pi. **B,** Microtubule pelleting assay of 1A_393_ and 1A-K1L12 in 2 mM ADP or ATP and BRB80. Plot is the percent of GFP-labelled motors in the microtubule pellet after centrifugation as a function of microtubule concentration. To account for any inactive motors, data are normalized to the concentration of motors bound in 1 mM AMPPNP, which induces a strongly-bound state of the motor. Fits with a Langmuir binding isotherm give the K_D_ and maximal binding fraction for each condition. 1A_393_ gives a Vmax of 97± 10 % and 102 ± 20 % in ADP and ATP, respectively. Despite the normalization to percent binding in AMPPNP, 1A-K1L12 does not reach full binding in this microtubule range with a Vmax of 24 ± 3 % and 26 ± 7 % in ADP and ATP, respectively. This could be because the K_D_ is actually around 30 μM or there is only ~50% maximal binding in ADP and ATP. Values are fit ± 95% CI and error bars are mean ± relative error. **C,** Dwell time distribution of 1A-K1L12 (light purple circles) in 2 mM ADP and BRB80. Biexponential fit gives dwell times of 0.03 ± 0.01 s and 0.3 ± 0.03 s (fit ± 95% confidence interval) with equal weights for the fast and slow populations. Data were collected at 50 frames per second. **D,** Dwell time distribution of Kin1 (pink circles) and K1-1AL12 (dark purple circles) in 2 mM ADP and BRB80. Values are single exponential fit ± 95% confidence interval. **E,** ATP-triggered Half-Site Release assay of 1A_368_ (black circles), 1A_393_ (blue circles), and 1A-K1L12 (light purple circles). Plot is the observed rate as a function of ATP concentration, and fitting with Michaelis-Menten equation give K_M_ and k_max_ values for each condition. For 1A_368_ (black circles), 1A_393_ (blue circles), and 1A-K1L12 (light purple circles), k_max_ were 172 ± 10, 154 ± 30, and 116 ± 13 s^-1^, respectively; K_M_ were 119 ± 20, 96 ± 53, and 64 ± 28 μM, respectively (fit ± 95% confidence interval).

To test whether the K-loop affects the detachment rate of KIF1A from its weak-binding state, we used an affinity assay in conjunction with a microtubule on-rate measurement. The dissociation constant is defined as 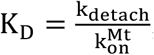; thus, by measuring both K_D_ and 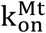 for each motor, we can calculate k_detach_. To estimate the K_D_ of 1A_393_ and 1A-K1L12, we carried out a microtubule pelleting assay in 2 mM ADP to mimic the weak-binding post-hydrolysis state. In this assay, varying concentrations of microtubules are mixed together with a constant concentration of motors, the mixture is pelleted, and the GFP fluorescence is used to determine the fraction of motors bound to the microtubule at each [Mt].^42^ As shown in Fig. 3B, swapping out Loop-12 of KIF1A reduced the microtubule affinity in ADP by four-fold, with K_D_ = 1.3 ± 0.8 μM for 1A_393_, and K_D_ = 5.1 ± 4.3 μM for 1A-K1L12. To relate these affinities to the affinity of the motor when it is processively walking along the microtubule, we repeated these pelleting assays in ATP. For both constructs, their K_D_ in ATP was similar to that in ADP (for 1A_393_, K_D_ = 1.7 ± 1.0 μM in ATP and for 1A-K1L12, K_D_ = 7.2 ± 5.1 μM in ATP; Fig. 3B), consistent with both motors spending most of their hydrolysis cycle in a weakly-bound ADP-like state, in agreement with previous work.^15^ To calculate the detachment rate from the weak-binding state, we used the K_D_ in ADP together with the 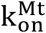 results from Fig. 2D, which were 10.6 ± 0.5 μM^-1^ s^-1^ for 1A_393_ and 9.1 ± 2.5 μM^-1^ s^-1^ for 1A-K1L12. From these values, we calculated k_detach_ in ADP to be 18 ± 11 s^-1^ for 1A_393_ and 66 ± 50 s^-1^ for 1A-K1L12, matching the 4-fold difference in the run lengths in Fig. 2B.

To more directly test whether the K-loop slows dissociation of KIF1A from the microtubule in the weakly-bound post-hydrolysis state, we used single-molecule TIRF microscopy to measure the dwell time of landing events in 2 mM ADP. For 1A_393_, the dwell time distribution was well fit by a single-exponential with a time constant of 1.8 ± 0.08 s (Fig. 1F), corresponding to an off-rate of 0.56 ± 0.02 s^-1^. When we repeated the experiment for 1A-K1L12, the kymographs showed a population of very transient events along with a population of longer duration events (Fig. S3). The dwell time distribution was well fit to a double exponential with a fast population of 0.03 ± 0.01 s (k_off_ = 33 s^-1^) that constituted ~40% of the events, and a slow population of 0.3 ± 0.03 s (k_off_ = 3.3 s^-1^) that constituted the other ~60 % of the events (Fig. 3C). Relative to the 1A_393_ dwell time of 1.8 s, these 1A-K1L12 dwell times correspond to a 60- and 6-fold faster off-rate when the K-loop of KIF1A is swapped out. Thus, both the microtubule pelleting assay and the single-molecule dwell time assay find that the apparent off-rate of KIF1A in the weakly-bound ADP state is increased when the K-loop is replaced by the equivalent sequence from kinesin-1.

As described by the kinetic race shown in Fig. 3A, it is also possible that the K-loop contributes to KIF1A processivity by enhancing the rate that the tethered head attaches to the microtubule during a forward step. To test this possibility, we carried out an ATP-triggered Half-Site Release assay, as follows (Fig. 3E).^49,50^ First, we incubated the motors with mant-ADP and microtubules to establish a complex of motors bound to the microtubule in a one-head-bound state, with mant-ADP trapped in the tethered-head. Then, we flushed this solution against varying concentrations of unlabeled ATP to initiate the binding of the tethered head and subsequent mant-ADP release. We next fit the fluorescent delay from the release of mant-ADP with an exponential function at each ATP concentration, plotted the observed rates versus the corresponding [ATP], and fit the data to a Michaelis-Menten curve. The observed rate constant at saturating ATP concentrations, k_max_, represents ATP hydrolysis, tethered head binding, and the subsequent release of mant-ADP. Because hydrolysis and mant-ADP are thought to be fast,^15^ k_max_ serves as a proxy for the tethered head binding rate. The k_max_ for our 1A_393_ construct (154 ± 30 s^-1^; fit ± 95% confidence interval) was similar to 1A_368_ (172 ± 10 s^-1^), indicating that the neck-coil does not alter the tethered head on-rate. For 1A-K1L12, k_max_ decreased slightly to 116 ± 13 s^-1^, indicating that, in addition to slowing the microtubule off-rate, the K-loop may enhance KIF1A superprocessivity by increasing the tethered head on-rate, 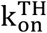. However, a caveat of this conclusion is that because the tethered-head attachment is thought to be the rate limiting step of the KIF1A chemomechanical cycle,^15^ a decrease in this rate should also decrease the overall motor velocity. Instead, the velocity for 1A-K1L12 was 20% faster than the 1A_393_ (Fig. 2C). Additionally, the calculated 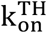 for 1A-K1L12 of 116 ± 13 s^-1^ is slower than the overall stepping rate of 212 s^-1^ (calculated as 1.7 μm/s velocity ÷ 8 per step). Thus, the question of whether the K-loop enhances the tethered-head on-rate is inconclusive from this experiment.

As a second strategy for estimating the tethered head attachment rate, we calculated it based on the observed run length in ATP and the measured detachment rate in ADP, as follows. From the kinetic race shown in Fig. 3A,^17^ the probability of detaching during each cycle is:

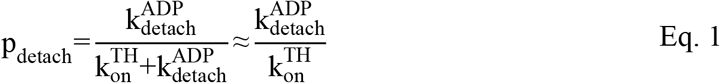

The number of steps a motor takes before dissociating can be estimated as the inverse of the detachment probability:

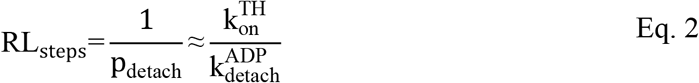

Thus, the tethered head attachment rate can be estimated by multiplying the measured detachment rate in ADP by the measured run length in ATP:

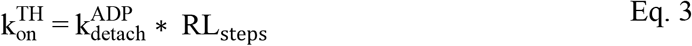

Table 1 shows the off-rates in ADP calculated from measured dwell times in Fig. 3 C and D, along with the run lengths calculated in number of steps from Fig. 2 B and E, and the resulting calculated tethered head attachment rate. The first result is that the calculated tethered head on-rates are roughly three-fold faster for KIF1A than kinesin-1, consistent with the faster stepping rate of KIF1A. The key result from this analysis is that for KIF1A, swapping out the K-loop has no effect on the calculated tethered-head on-rate. In summary, the K-loop contributes to the superprocessivity of KIF1A by slowing the off-rate of the motor from the vulnerable 1HB state, and the K-loop does not modulate the processivity of KIF1A by enhancing the tethered head attachment rate.

**Table 1:**
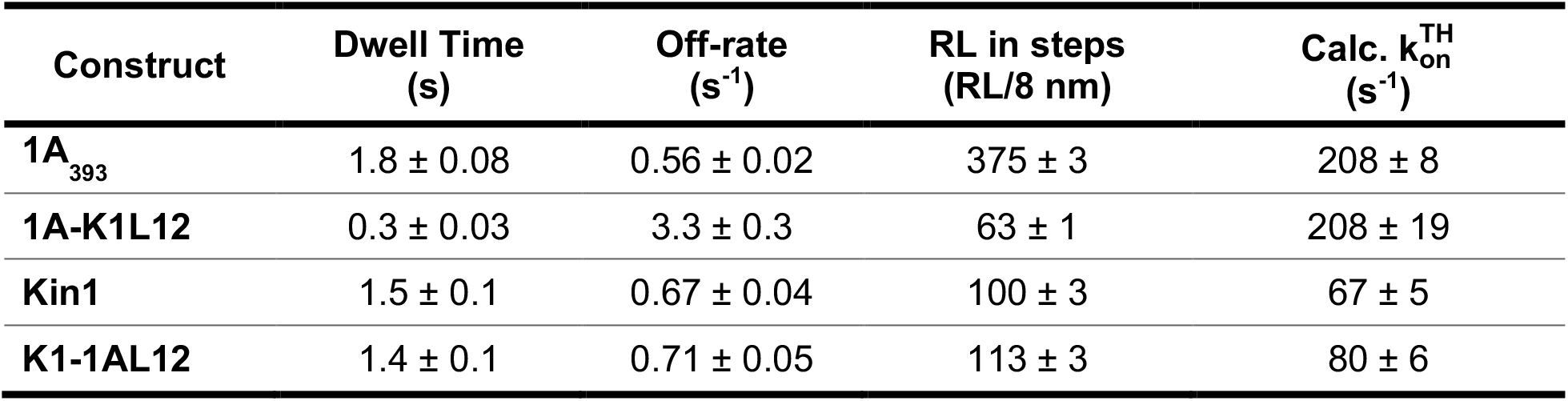
Calculations of Tethered-head attachment rate for different constructs. Dwell times are fit ± 95% CI from Fig. 3 and off-rates are inverse of dwell times with propagated errors. Run Length (RL) in steps are the RL taken from Fig. 2 divided by the step size (8-nm), and errors are ± 95% CI. 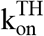 were calculated using Eq. 3, with errors propagated from off-rates and RL.

### KIF1A run length scales with charge of the K-loop

In principle, swapping in the K-loop of kinesin-1 could be reducing the KIF1A run length either solely due to differences in positive charge, or through some combination of charge and the length of the loop. To test whether the run length data can be accounted for exclusively based on the charge of loop-12, we designed three additional loop-12 mutant constructs having the same length as wild type but having varying net charge in the loop-12 domain. First, we increased the total charge of loop-12 in our 1A_393_ construct by replacing three of the native uncharged residues with lysines, resulting in a net charge of +7 in loop-12; we refer to this construct as SuperK (Fig. 4A). Using single-molecule motility assays in BRB80 at 2 mM ATP, we found that the run length of SuperK was ~2-fold longer and the segmented velocity was slightly slower compared to the control 1A_393_ (Fig. 4B-C; Table 2). We also used stopped-flow to measure the bimolecular on-rate, 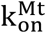, and found that the SuperK on-rate was 1.5-fold faster than 1A_393_, but the values were within fit error of one another (Fig. 4D).

**Figure 4:**
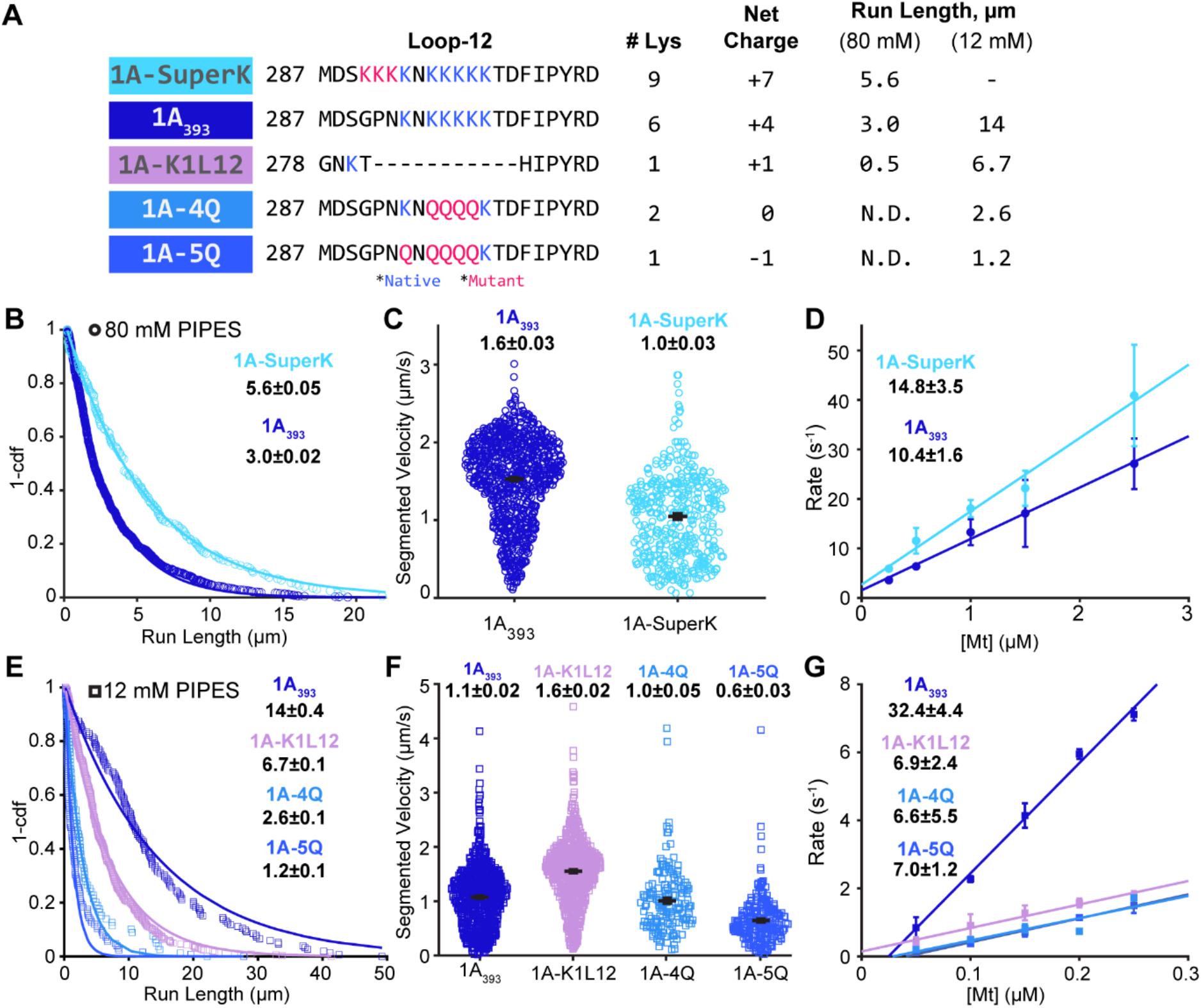
Charge of the KIF1A K-Loop regulates superprocessivity. **A,** Sequence of the Loop-12 domain in the protein constructs used, along with the number of lysine residues and net charge of the domain. Mutations are in red; native lysines are in green. ND indicates not detectable run length. Representative kymographs in Fig. S4. **B,** Run length distribution of 1A_393_ (blue circles) and 1A-SuperK (light blue circles) in BRB80 buffer and 2 mM ATP. Values are single exponential fits ± 95% confidence interval. **C,** Velocity distribution of 1A_393_ (blue circles) and 1A-SuperK (light blue circles) in BRB80 and 2 mM ATP. Average velocities for segments with Δx > 3 pixels are mean ± SEM. **D,** Bimolecular on-rates of 1A_393_ (blue circles) and 1A-SuperK (light blue circles) in BRB80. Linear fits, reported as fit ± 95% confidence interval, give 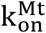. Points are mean ± SEM. **E,** Run length distribution of 1A_393_ (blue squares), 1A-4Q (light blue squares), 5Q (medium blue squares) and 1A-K1L12 (light purple squares) in BRB12 buffer and 2 mM ATP. Values are single exponential fit ± 95% confidence interval. **F,** Velocity distribution of 1A_393_ (blue squares), 1A-K1L12 (light purple squares), 1A-4Q (light blue squares), and 1A-5Q (medium blue squares) in BRB12 and 2 mM ATP. Average velocities for segments with Δx > 3 pixels are reported as mean ± SEM. **G,** Bimolecular on-rates of 1A_393_ (blue squares), 1A-4Q (light blue squares), 1A-5Q (medium blue squares) and 1A-K1L12 (light purple squares) in BRB12 buffer. Linear fits give 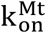 (fit ± 95% confidence interval). Points are mean ± SEM.

**Table 2:** Summary of data in BRB80 buffer.

| Construct | RL ($\mu\text{m}$) | Vel ($\mu\text{m/s}$) | ADP Dwell (s) | $k_{on}^{Mt}$ ($\mu\text{M}^{-1} \text{s}^{-1}$) |
| --- | --- | --- | --- | --- |
| **1A(1-393)** | $3.0 \pm 0.02$ | $1.6 \pm 0.03$ * | $1.8 \pm 0.08$ | $10.6 \pm 0.5$ |
| **1A-K1L12** | $0.5 \pm 0.004$ | $1.7 \pm 0.03$ | $0.3 \pm 0.03$ | $9.1 \pm 2.5$ |
| **K1** | $0.8 \pm 0.02$ | $0.7 \pm 0.01$ | $1.5 \pm 0.1$ | $1.7 \pm 0.6$ |
| **K1-1AL12** | $0.9 \pm 0.02$ | $0.8 \pm 0.01$ | $1.4 \pm 0.1$ | $9.9 \pm 2.9$ |
| **Super-K** | $5.6 \pm 0.05$ | $1.0 \pm 0.03$ * | - | $14.8 \pm 3.5$ |

\*Segmented Velocity reported.

Next, to test whether reducing the net charge of loop-12 reduces the run length, we replaced a portion of the of the lysine residues in loop-12 with glutamines. Glutamine was chosen because it is uncharged in our buffer (pH 6.9) and the side chain is of similar size to lysine, minimizing potential steric effects. We substituted four or five lysines in 1A_393_ by glutamines, creating 4Q and 5Q, which had net charges in Loop-12 of 0 and −1, respectively (Fig. 4A). In BRB80 buffer, we observed no processive events for 4Q or 5Q (Fig. 4A; ND, not detected). Thus, although swapping in the kinesin-1 K-loop (K1L12; +1 net charge) led to short but detectable processivity, reducing the net charge of the K-loop further led to undetectable processivity in BRB80 buffer.

Because lowering the ionic strength enhanced the run length of other KIF1A constructs, we tested the processivity of these K-loop charge mutants in BRB12 buffer and found that they had measurable run lengths under these conditions (see representative kymographs in Fig. S4). This result confirms that the lack of events in BRB80 was not due to protein misfolding or other off-target effects. In BRB12, the run lengths of 4Q and 5Q scaled with the net charge of loop-12, as follows (Fig. 4E). Wild-type 1A_393_ (+4 charge) had a run length 14 ± 0.4 μm; K1L12 (+1 charge) had a run length of 6.7 ± 0.1 μm; 4Q (neutral) had a run length of 2.6 ± 0.1 μm; and 5Q (−1 net charge) had a run length of 1.2 ± 0.1 μm) (see also Fig. 6 and Table 3). The velocities in BRB12 also differed, but not in a charge-dependent manner (Fig. 4F; Table 3). Interestingly the bimolecular on-rates of the three mutants in BRB12 buffer were similar to one another and they were all ~5-fold slower than 1A_393_ (Fig. 4G). Thus, the key feature of KIF1A loop-12 that enhances the motor’s processivity is the positive charge rather than the longer length of the loop relative to kinesin-1.

**Figure 5:**
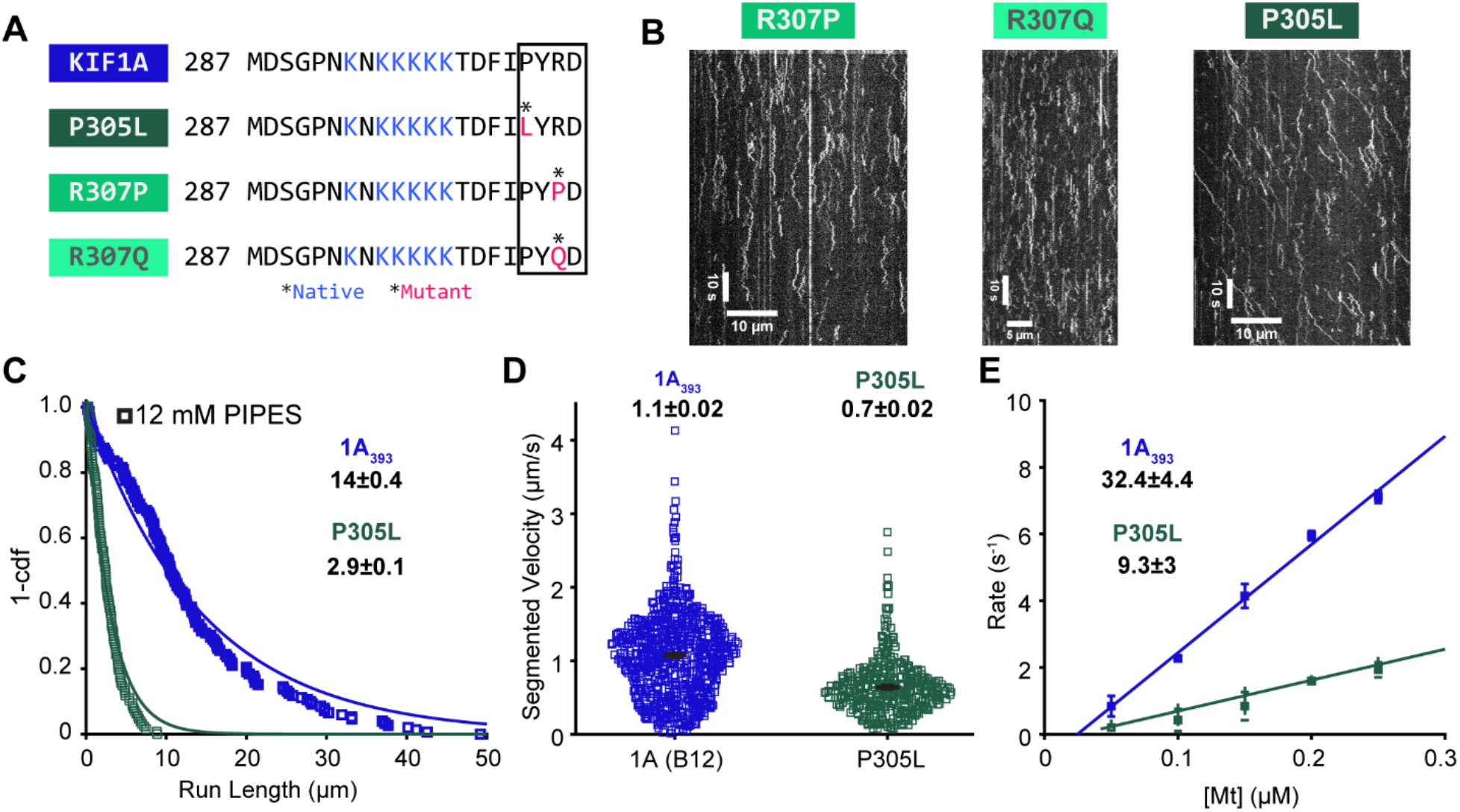
Pathogenic mutations in Loop-12 hinder KIF1A motility. **A,** Sequence of the Loop-12 for wild type and mutants, with point mutations indicated in red and conserved distal region indicated by a box. **B,** Example kymographs of R307P, R307Q, and P305L in BRB12 buffer and 2 mM ATP. **C,** Run length distribution of 1A_393_ (blue squares) and P305L (green squares) in BRB12 and 2 mM ATP. Values are single exponential fit ± 95% confidence interval. **D,** Velocity distribution of 1A_393_ (blue squares) and P305L (green squares) in BRB12 and 2 mM ATP. Average velocities for segments with Δx > 3 pixels are mean ± SEM. **E,** Bimolecular on-rates of 1A_393_ (blue squares) and P305L (green squares) in BRB12. Linear fits give 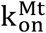 (fit ± 95% confidence interval). Points are mean ± SEM.

**Figure 6:**
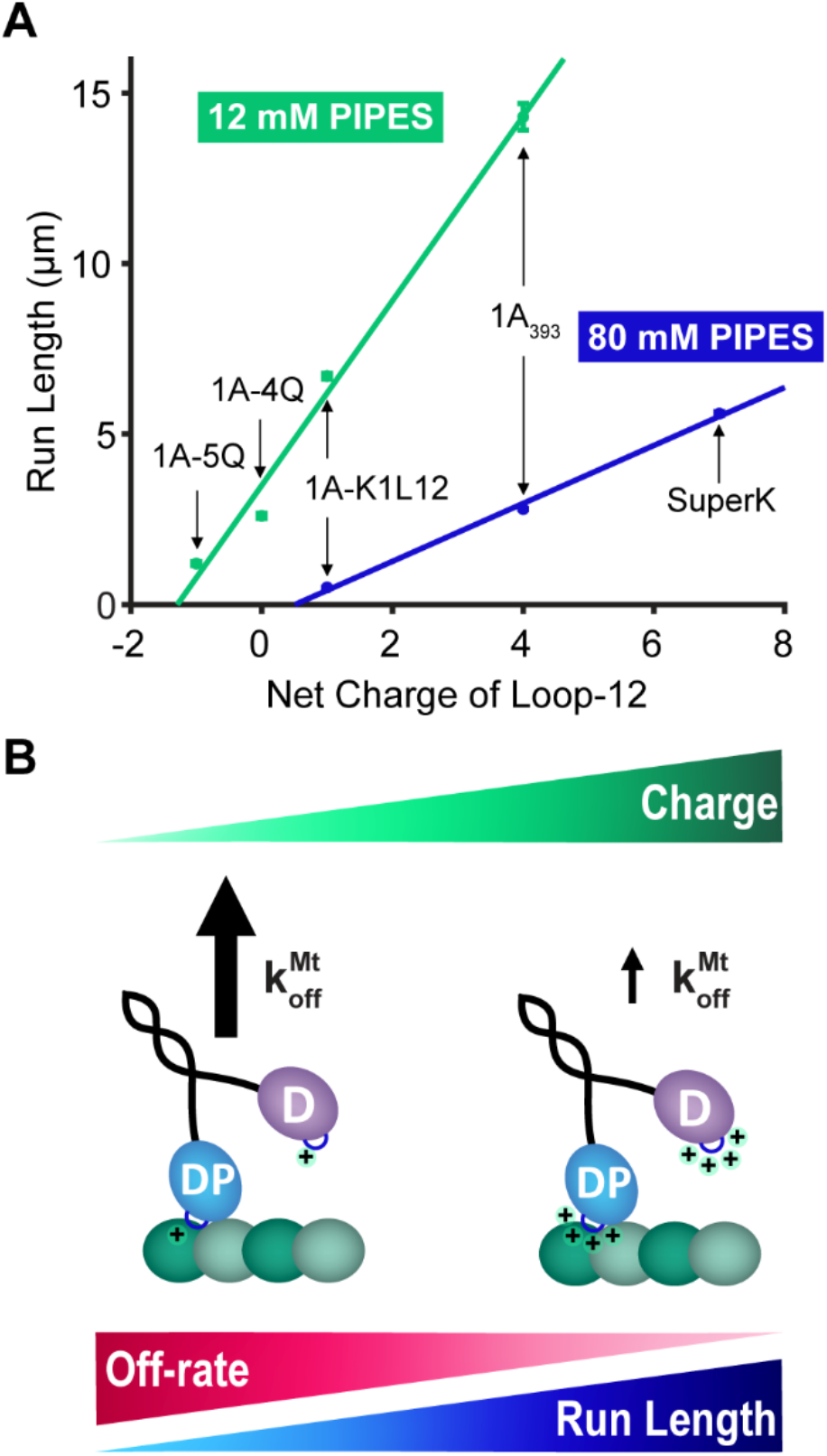
Run Length of KIF1A is dependent on Charge of Loop-12. **A,** Run Length of the various processive constructs used in this study, plotted as a function of the net charge of their Loop-12 domain, in BRB12 and BRB80. Points are from the 1-CDF plots in previous figures (fit ± 95% CI). Linear fits to the points in BRB12 and BRB80 give slopes of 2.7 ± 0.9 and 0.9 ± 0.6, respectively (fit ± 95% CI). **B,** Diagram depicting how increasing positive charge in Loop-12 leads to a slower off-rate in ADP that results in longer run lengths.

**Table 3:** Summary of Data in BRB12 buffer.

| Construct | RL ($\mu\text{m}$) | Vel ($\mu\text{m/s}$) | $k_{on}^{Mt}$ ($\mu\text{M}^{-1} \text{s}^{-1}$) |
| --- | --- | --- | --- |
| **1A(1-393)** | $14.3 \pm 0.4$ | $1.1 \pm 0.02$ * | $32 \pm 4.4$ |
| **1A-K1L12** | $6.7 \pm 0.1$ | $1.6 \pm 0.02$ * | $6.9 \pm 2.4$ |
| **4Q** | $2.6 \pm 0.1$ | $1.0 \pm 0.05$ * | $6.6 \pm 5.5$ |
| **5Q** | $1.2 \pm 0.1$ | $0.6 \pm 0.03$ * | $7.0 \pm 1.2$ |
| **P305L** | $2.9 \pm 0.1$ | $0.7 \pm 0.02$ * | $9.3 \pm 3$ |

\*Segmented Velocity reported.

### Human disease mutations adjacent to the K-loop impair KIF1A motility

To extend our understanding of the role of loop-12 in KIF1A motility, we investigated the motile properties of a set of mutations identified in patients suffering from KIF1A Associated Neurological Disorder (KAND). A recent study investigated the clinical features of 117 KAND patients and identified a number of mutations in the well conserved ‘PYRD’ sequence at the carboxyl end of loop-12; no patients in the study had mutations in the K-loop proper.^14^ We investigated, three pathogenic human mutations: R307P, R307Q, and P305L (Fig. 5A). The four patients harboring the R307Q mutation had moderate to severe KAND with hypertonia/spasticity, and the pair of twins that harbored the R307P mutation displayed brain atrophy and seizures.^13,14^ In *C. elegans,* R307Q was able to partially rescue a null mutant;^51^ however, to our knowledge the single-molecule properties of R307Q and R307P have not been evaluated. The four patients harboring the P305L mutation ranged from mild to severe KAND; all showed hypertonia/spasticity, but only two of four showed brain atrophy and one of the four exhibited seizures. In contrast to R307Q, P305L was unable to rescue a *C. elegans* null mutant; however, P305L was shown to be motile in single-molecule assays albeit with impaired performance.^52^ Thus the P307 mutants, which decrease positive charge near the K-loop, present more severe clinical phenotypes, but the P307Q can partially rescue worms. In contrast, P305L, which has been proposed to alter the conformation of a helix adjacent to the K-loop, has a less severe clinical phenotype and the motor retains some motility, but is unable to rescue mutant worms.^52^

We first examined R307P, R307Q, and P305L mutants using single-molecule tracking in BRB80 and 2 mM ATP, and observed no motility for any of the three disease mutants (Fig. 5B). When the ionic strength was lowered using BRB12 buffer, R307P and R307Q showed a higher frequency of landing events and longer duration of diffusive events, but still no persistent directional movement (Fig. 5B). In contrast, P305L did show processive movement in BRB12, but with a ~5-fold shorter run length and ~2-fold slower velocity than the 1A_393_ control in BRB12 (Fig. 5C-D). A previous study found that that the P305L mutation strongly reduced the microtubule landing rate, and suggested that the mutation alters the interaction of the K-Loop with the C-terminal tail of tubulin.^52^ To directly measure the microtubule on-rate of P305L, we used the stopped flow 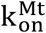 assay in BRB12 and found that the P305L mutant had a ~3-fold lower on-rate than 1A_393_ (Fig. 5E). Interestingly, the P305L on-rate of 9.3 ± 3 μM^-1^s^-1^ (Fig. 5E) was faster than either the loop swap mutant 1A-K1L12 or the two glutamine mutants in BRB12 (Fig. 4G; Table 3). This ~3-fold decrease in the on-rate was in the same direction, but it was much smaller than the ~35-fold decrease in the microtubule landing rate of P305L in previous work.^52^

## Discussion

The positively charged loop-12 in kinesin-3 motors, known as the K-Loop, has been a topic of interest in the field for many years, but there has yet to be a consensus on the role of the K-loop in the mechanism of KIF1A superprocessivity. In this work, we have established a comprehensive understanding of the mechanism of the K-Loop in the KIF1A chemomechanical cycle. We conclude that the unique superprocessivity of KIF1A dimers originates from the charge-dependent interaction of loop-12 with the microtubule, resulting in a reduction in off-rate from the post-hydrolysis one-head-bound (1HB) state (Fig. 6). This stabilization of the weak-binding state allows time for the tethered head to bind to the next binding site and complete the step, and thus maximizes the number of steps the motor takes before terminating a processive run.

Despite an appreciation of the importance of the K-loop, there are contradictions in the literature regarding the role of the K-loop in KIF1A superprocessivity. Early work made the striking finding that KIF1A monomer constructs move processively through a combination of intermittent forward steps and 1D diffusion along the microtubule.^25,53^ The microtubule affinity of these monomers was shown to depend on the negatively charged C-terminal tail of tubulin, scale with the amount of positive charge in loop-12, and be enhanced in low ionic strength buffers. Subsequently, it was shown that full-length KIF1A is dimeric and the motility of dimer constructs was fast, superprocessive, and lacked the diffusional behavior of the monomers.^30,54^ However, using a KIF1A dimer truncated after the neck-coil (1-393 aa), it was found that replacing the KIF1A loop-12 with that of kinesin-1 did not diminish the run length,^31^ a result that was puzzling in light of the monomer results. Subsequent work showed that, unlike kinesin-1, the neck-coil domain of KIF1A is insufficient to stably dimerize the motor, and stabilized dimers could be created by adding a leucine zipper sequence downstream of the neck-coil.^19,27,30,32^ Upon closer inspection, the apparent lack of influence of loop-12 on KIF1A processivity can be explained by the fact that the construct used in that work (KIF1A(1-393)) lacked a distal coiled-coil region that is needed to stabilize the dimer.^31^ In that work, KIF1A(1-393) had a run length of 2.6 μm, the loop-swap mutant had a run length of 3.6 μm, and the stably dimerized KIF1A(1-393)-LZ had a run length of 9.8 μm.^31,32^ Thus, the apparent lack of influence of the K-loop in the truncated dimer lacking a stabilizing LZ domain can be explained by run lengths being terminated by the motor reverting to monomers and dissociating from the microtubule rather than termination of a processive run by normal dissociation of the dimer. As a competing process, this premature termination would mask any change in run length due to the loop swap.

By measuring key transitions in the kinetic race that defines processivity, we find that the principal role of the K-loop is to stabilize the weak-binding, post-hydrolysis state of KIF1A. The finding from the pelleting assay that the K_D_ for microtubules is similar in ADP and ATP for both wild-type and Loop-12 swapped KIF1A emphasizes the importance of the weak binding state for KIF1A processivity. Their similar residence times are consistent with KIF1A spending the majority of its cycle in a vulnerable one-head-bound ADP state, a key finding from a previous study.^15^ The parallel reduction of the microtubule affinity in ADP and ATP upon swapping out the K-loop clearly demonstrates how decreasing the microtubule affinity in the ADP state dictates a reduction in run length. Because the other component of the kinetic race, tethered head attachment, also involves microtubule binding and hence might be expected to be modulated by charge in the K-loop, it was notable that swapping out the K-loop did not alter the tethered head on-rate. Tethered head binding involves both a mechanical component of the tethered head stretching to the next binding site, as well as a biochemical component of ADP release to achieve tight binding, and so the simplest interpretation of this result is that the tethered head binding rate is not substantially mediated by electrostatic interactions between the motor domain and the microtubule. Surprisingly, we found that swapping out the K-loop of KIF1A did not diminish the bimolecular on-rate in solution, meaning that electrostatics mediated by the K-loop do not play a strong role in the initial landing of KIF1A on the microtubule at near physiological ionic strength. This behavior contrasts with kinesin-1, where inserting the K-loop of KIF1A did enhance the bimolecular on-rate in solution, a difference that highlights the different kinetic tuning of the two motor families involved in the initial binding of the motor to the microtubule.

By systematically replacing lysines in KIF1A loop-12 with glutamines, we find that the run length of KIF1A scales linearly with the net charge of loop-12 in both BRB80 and BRB12 (Fig. 6). Three published studies using KIF1A or the C. elegans ortholog Unc104 in BRB12 buffer found similarly that reducing the charge of the K-loop by either swapping with the kinesin-1 sequence or replacing lysines with alanines decreased the run length.^27,30,31^ Arpag *et. al.* found that swapping the KIF1A K-loop (net charge +4) with the rat kinesin-1 sequence (net charge +1) caused a 2.2-fold reduction in run length, and Lessard found a 2.6-fold reduction in run length when the net charge of the K-loop was reduced from +4 to +1 by replacing three lysines in the K-loop with alanines. ^27^ These values are similar to the 2.1-fold reduction in run length we measured between 1A_393_ (net charge +4) and 1AK1L12 (net charge +1) (Fig. 6A). Using the *C. elegans* KIF1A ortholog Unc104, Tomishige found that swapping the K-loop (net charge +4) with the human kinesin-1 sequence (net charge 0) resulted in a 5.4-fold reduction in run length, which matches our 5.5-fold shorter run length for K1-4Q (net charge 0). The similarity across these studies (shown graphically in Fig. S5) reinforces our finding that KIF1A processivity scales linearly with the charge of the K-loop, and also argues that this reduction does not depend on the specific sequences, but solely due to the electrostatic charge. This linear relationship and x-intercept around a −1 charge also helps to explain why a construct in which all six lysines in the K-loop were replaced with alanines, resulting in a net charge of −2, was not measurably processive.^27,31^ Finally, the shallower slope and more positive x-intercept in BRB80 in Fig. 6A is consistent with charge shielding at higher ionic strength reducing the impact of electrostatic interactions on the run length.

We also found that in near-physiological ionic strength buffer, swapping the KIF1A K-loop into kinesin-1 did not confer superprocessivity on this motor (Fig. 2). This result suggests that the KIF1A chemomechanical cycle is tuned such that it relies on the K-loop to achieve superprocessivity, whereas the kinesin-1 chemomechanical cycle is tuned to rely on different mechanisms to achieve processivity. One potential explanation is that because kinesin-1 spends a much smaller fraction of its ATP hydrolysis cycle in a one-head bound vulnerable state than KIF1A,^15,36^ altering the microtubule affinity of this state has a negligible effect on the run length. However, the microtubule detachment rate of kinesin-1 in the weak-binding ADP state was not altered by swapping in the KIF1A K-loop (Fig. 3D), arguing against this mechanism and suggesting instead that the positive charge in the K-loop cannot stabilize this vulnerable state in kinesin-1 the way it can in KIF1A. One possibility is that during transient episodes when the motor domain is tethered to the microtubule solely by its K-loop, KIF1A rebinds rapidly through its canonical microtubule binding site to maintain association, whereas kinesin-1 rebinds more slowly and instead dissociates from this tethered state despite the added electrostatic interactions. This difference in weak-binding characteristics suggests that binding of kinesin-1 to the microtubule in the ADP state is dominated by a different region of the microtubule binding domain, such as Loop-8 or Loop-11/a-4.^55^

There are a number of documented KAND mutations in the well-conserved ‘PYRD’ sequence at the carboxyl end of loop-12, ^14^ but how these mutations alter the chemomechanical cycle or the interaction of KIF1A with the microtubule is not clear. Despite a recent report that an R307Q mutant partially rescues vesicle transport in a null-mutant worm,^51^ we found that R307Q and R307Q were incapable of productive movement in either BRB80 or BRB12 (Fig. 5). Published molecular dynamics simulations found that R307 in KIF1A (and the equivalent R278 in kinesin-1) interact electrostatically with residues in the tubulin core, and thus likely contribute to the strength of microtubule binding.^55,56^ The calculated binding free energies between R307 and residues in tubulin were similar between the strong-binding ATP and Apo states and the weak-binding ADP state in that work,^55^ suggesting that R307 is not involved in nucleotide-dependent changes in microtubule binding affinity that occur through the KIF1A mechanochemical cycle. However, the lack of motility of the R307 mutants suggests this residue plays a key role in mechanochemical coupling in the motor domain and/or the strong binding interaction needed for stepping. On the other hand, the diffusive binding of both R307P and R307Q suggest that these mutations do not prevent the K-loop from interacting with the C-terminal tail of tubulin. A nearby mutation, P305L, was previously proposed to alter the interaction of the K-loop with the microtubule based on a ~35-fold reduction in the single-molecule landing rate.^52^ That work was carried out in a ~160 mM ionic strength HEPES buffer at pH 7.4, though a different study using ~80 mM ionic strength HEPES buffer at pH 7.2 found only a two-fold decrease in the landing rate. ^14^ Using a biochemical assay in BRB12 (26 mM ionic strength, pH 6.9) that is robust to variations in motor activity between constructs, we found a 3.5-fold decrease in the bimolecular on-rate for P305L. This on-rate suppression was smaller than the effect of mutations that neutralized the charge of the K-loop (see Table 3), which suggests that at low ionic strength the P305L mutation does not inhibit the interaction of the K-loop with the C-terminal tail of tubulin. Notably, we failed to observe any motility of P305L in BRB80 (173 mM ionic strength, pH 6.9), which is consistent with the inability of P305L to rescue vesicle motility in a null worm.^52^ Hence, the effect of different KAND mutations on microtubule binding and chemomechanical coupling are highly dependent on experimental conditions, and further work is needed to connect the structural changes with alterations in motility, as well as how changes in the motile properties translate to defects in axonal transport.

The present study highlights different chemomechanical tuning strategies that kinesin-1 and kinesin-3 employ to carry out their intracellular transport functions. KIF1A is notable in being fast and superprocessive, and it does this by spending most of its hydrolysis cycle in a vulnerable one-head-bound state that is stabilized by electrostatic interactions between the K-loop and the microtubule. Importantly, this strategy causes the motor to detach readily under load, which is seemingly not an advantageous property for a transport motor.^19–22^ However, KIF1A binds to the microtubule from solution at a much faster rate than kinesin-1, which may partly compensate for this detachment.^15^ In contrast, kinesin-1 is able to walk processively against substantial loads,^20,22^ but it walks more slowly, it binds to the microtubule out of solution more slowly, and in the absence of load has a considerably shorter run length.^15,31^ As the mechanistic features of these motors become clarified, the next step is to understand how these functional properties scale up to multimotor cargo transport on diverse microtubules and bidirectional tug-of-war transport with dynein. Similarly, to understand how specific mutations lead to KAND disease states, it will be necessary to define the effect of other mutations on KIF1A chemomechanics and to extrapolate how those changes affect neuronal function.

## Methods

### Protein Preparation

*Rattus norvegicus* KIF1A motor constructs were based on previous work.^15^ The 1A_393_ construct, used as the ‘wild type’ throughout this study, included the KIF1A motor head, neck linker, and neck-coil domains (residues 1-393) followed by the *D. melanogaster* kinesin-1 neck-coil and coiled-coil 1 domains (residues 345 to 560), and C-terminal GFP tag for single-molecule assays, or *D. melanogaster* kinesin-1 neck-coil (residues 345 to 406) for biochemical assays. The 1A_368_ construct consisted of the KIF1A motor head and the neck linker domains (residues 1-368) followed by the *D. melanogaster* kinesin-1 coil domains for stable dimerization (same residues and tags as used above for 1A_393_). The 1A-LZ construct was composed of *R. norvegicus* KIF1A residues 1-393 followed by a leucine zipper domain (see Fig. S2 for sequence) for stable dimerization and a C-terminal GFP tag.^32^ Subsequent constructs used throughout this study introduced mutations (as described in the Results) using the 1A_393_ construct as the template. All proteins contained a C-terminal His6 tag. Plasmids were designed in SnapGene and mutants were generated via Gibson assembly (Gibson Assembly® Cloning Kit from New England Biolabs)^57,58^ or via QuickChange (Q5® Site-Directed Mutagenesis Kit from New England Biolabs).^59^ Recombinant expression of motors in *E. coli* and purification via Ni-NTA chromatography were performed as described previously.^15^

### Single Molecule Fluorescence Tracking

Single-molecule TIRF microscopy assays were performed as previously described.^15,40,42^ Flow cells were injected sequentially with casein for surface blocking, rigor motors for binding microtubules to the surface,^36^ and taxol-stabilized microtubules in BRB80 (80 mM PIPES, 1 mM EGTA, 1 mM MgCl_2_, pH 6.9). The imaging buffer mixture was BRB80 with 20 mM D-glucose, 0.02 mg/ml glucose oxidase, 0.008 mg/ml catalase, and 10 mM DTT (dithiothreitol). Motors were diluted in imaging buffer with 0.5 mg/ml casein and 2 mM ADP/ATP (and equal molar MgCl_2_). Videos were recorded at 10 frames per second in most cases, except 20 or 50 fps in ADP and BRB80 and 5 fps in ADP or ATP and BRB12. Kymograph analysis was carried out manually with ImageJ. To eliminate bias, all traces below 3 pixels (along the distance axis for ATP assays and the time axis for ADP assays) were excluded from the data set. Fits to the data distributions were done using MATLAB R2020b (MathWorks). The cutoff for the fits in the 1-CDF plots was determined as 3x the pixel value (i.e. 0.17 μm for run length in ATP assays or 0.3 s for ADP dwell time assays done at 10 fps). Statistical analysis was carried out using GraphPad Prism7 t-test function (Kolmogorov-Smirnov test) to determine significance relative to the control.

### Stopped Flow Assays

Stopped Flow assays, including k_on_^Mt^ and ATP-triggered Half-Site Release were performed as previously described.^15^ Concentrations mentioned below refer to the ‘syringe concentrations’ and the final concentrations in the chamber were half. The k_on_^Mt^ assays in BRB80 were performed by flushing 300 nM of active motor dimers in 0.5 μM mant-ADP against 2 mM Mg-ADP and varying concentrations of taxol-stabilized microtubules (0.5 - 5 μM). In BRB12, 300 nM of active motor dimers in 5 μM mant-ADP were flushed against 2 mM Mg-ATP and varying concentrations of taxol-stabilized microtubules (0.1 – 0.5 μM). The ATP-triggered Half-Site Release assays were performed by incubating 0.5 μM active motor dimers with 6 μM of taxol-stabilized microtubules and 1 μM of excess mant-ADP to form a one-head-bound complex in solution. This complex was then flushed against varied concentrations of Mg-ATP.

### Microtubule Pelleting Assays

In microtubule pelleting assays, GFP-labeled motors were incubated with varying concentrations of taxol-stabilized microtubules with 2 mM ADP or ATP. After a 5 minute incubation at room temperature, the solutions were spun down using an Airfuge at 30 psi for 10 minutes. A Molecular Devices FlexStation 3 Multimode Microplate Reader was used to measure the GFP concentration of the pellet and supernatant. Reported values (% pellet) were calculated as the fluorescence of the pellet divided by the sum of the pellet and supernatant. All values were normalized to a control value of motors in 2 mM AMPPNP and 20 μM taxol-stabilized microtubules.

## Acknowledgements

The authors thank Dr. Kristen Verhey for helpful discussions that resolved differences between the current results and previously published KIF1A data. The authors thank the Hancock lab group members for their helpful discussions, with particular thanks to Adheshwari Ramesh and Samira Bell for their help making the plasmids used in this study.

## Supplemental Information

**Figure S1:**
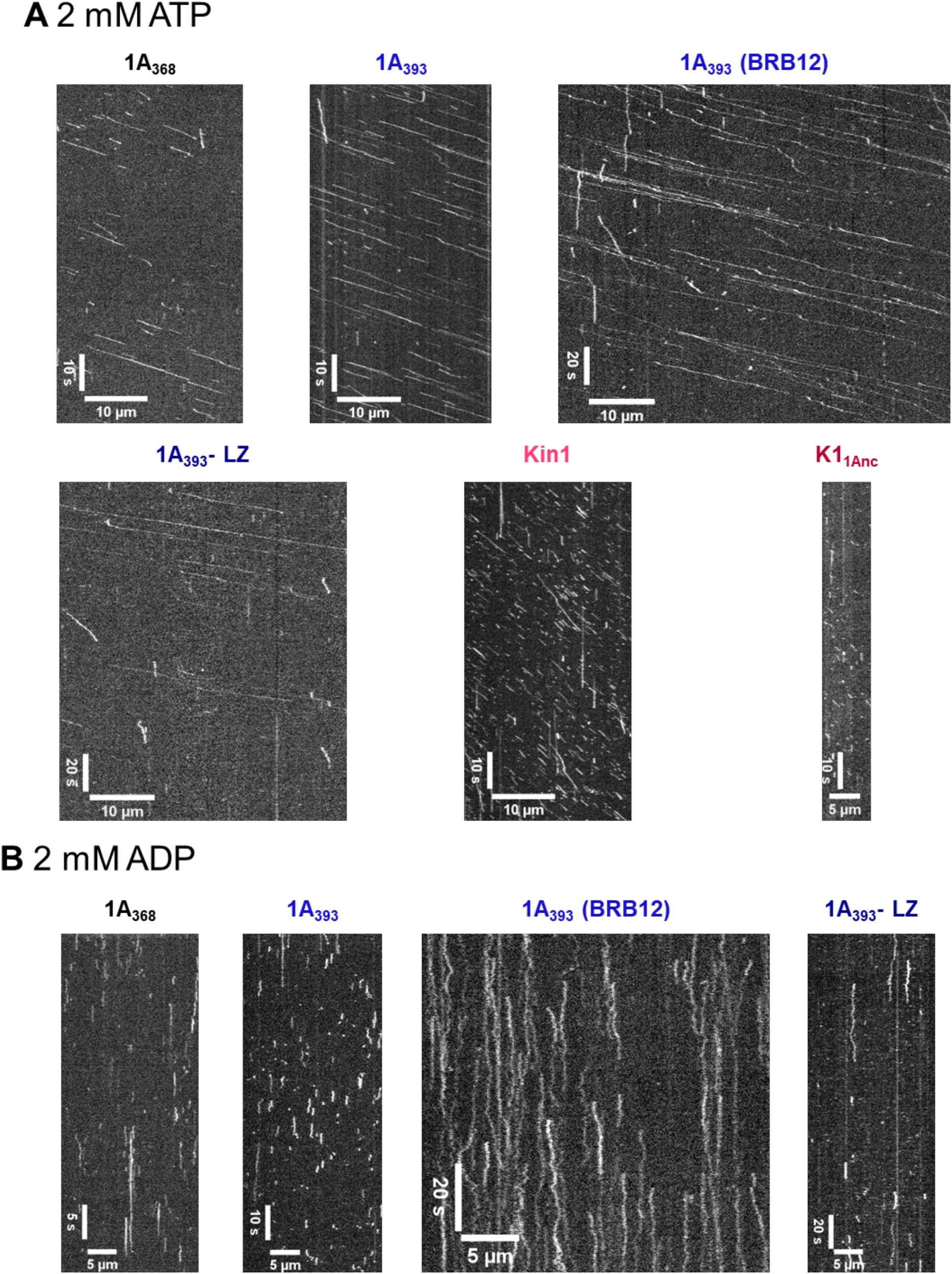
Example kymographs for different KIF1A and Kin1 constructs in ATP and ADP. **A,** Kymograph of each construct in 2 mM ATP and BRB80 (unless otherwise noted). **B,** Kymograph of each construct in 2 mM ADP and BRB80 (unless otherwise noted). Individual kymographs were analyzed from videos at various frame rates; scale bars reflect this distinction.

**Figure S2:**
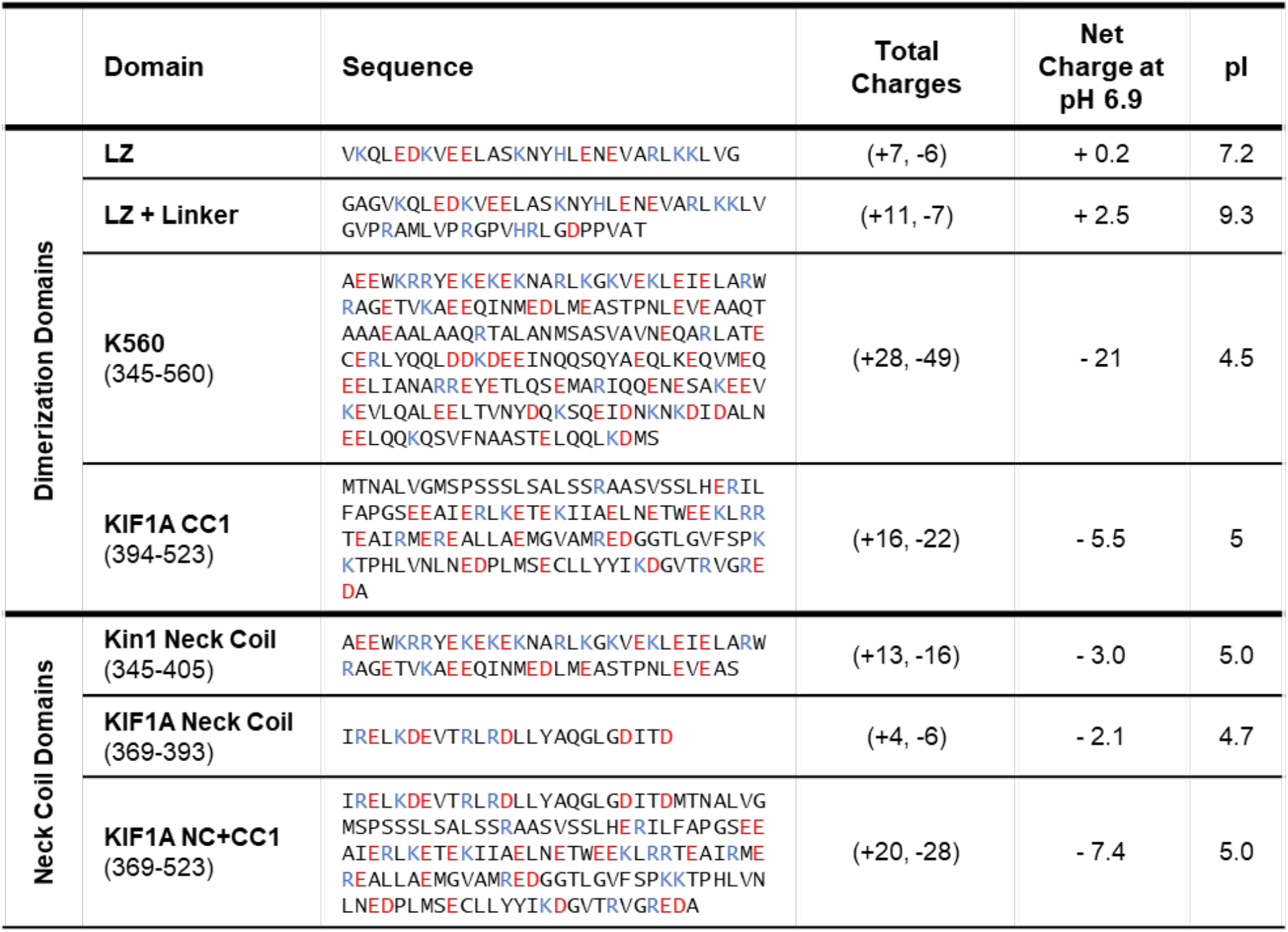
Charge of the neck coil domains impacts motor properties of KIF1A. **A,** Table of the different dimerization and neck coil domains used in this study with the sequence, total number of charged residues, net charge of the domain at pH 6.9 and the pI. Net charge and pI were calculated using (http://protcalc.sourceforge.net/)

**Figure S3:**
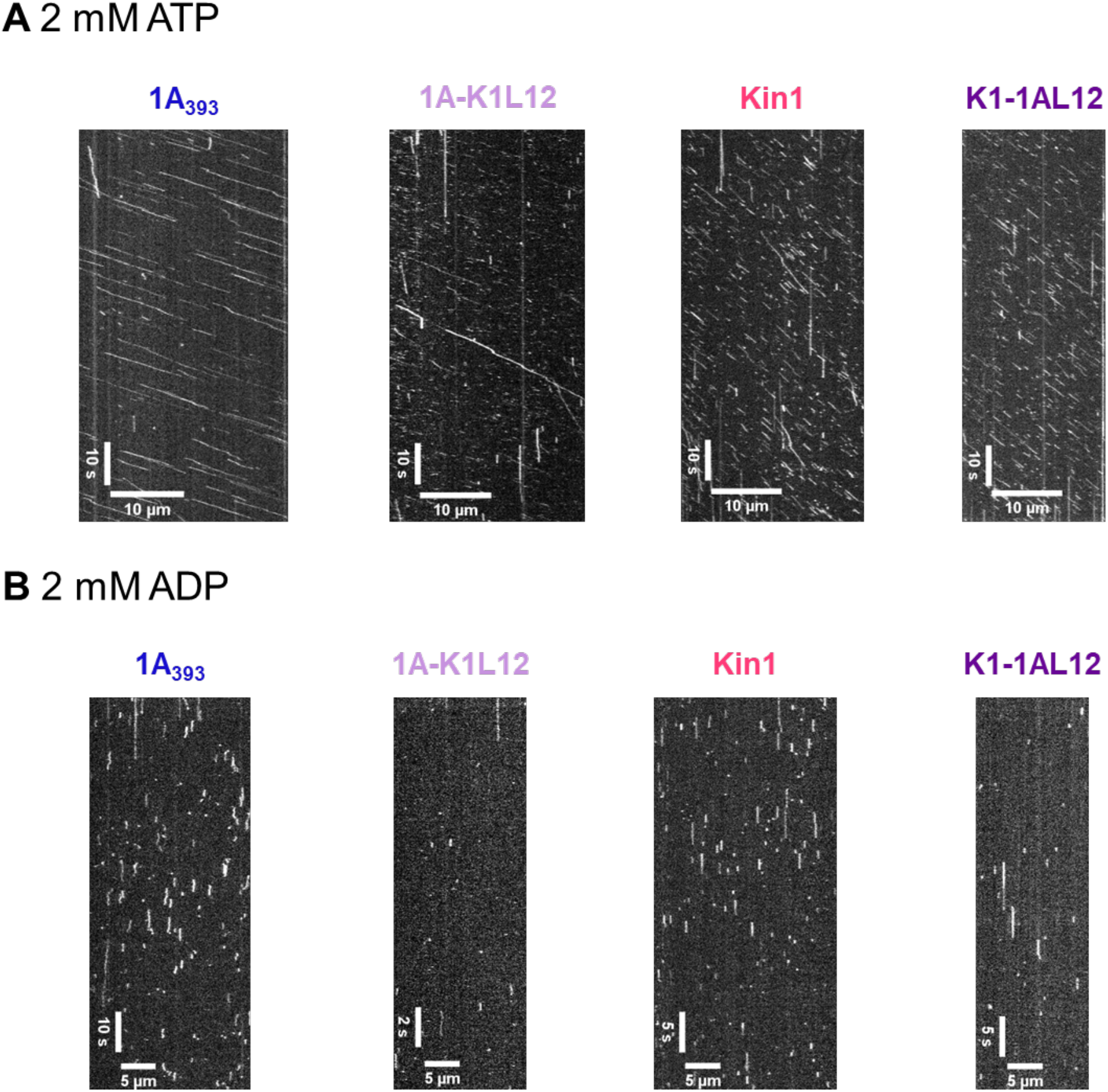
Example kymographs of constructs in ATP and ADP. **A**, Kymographs of different constructs used in Fig. 2, in 2 mM ATP and BRB80. **B**, Kymographs of different constructs used in Fig. 2, in 2 mM ADP and BRB80.

**Figure S4:**
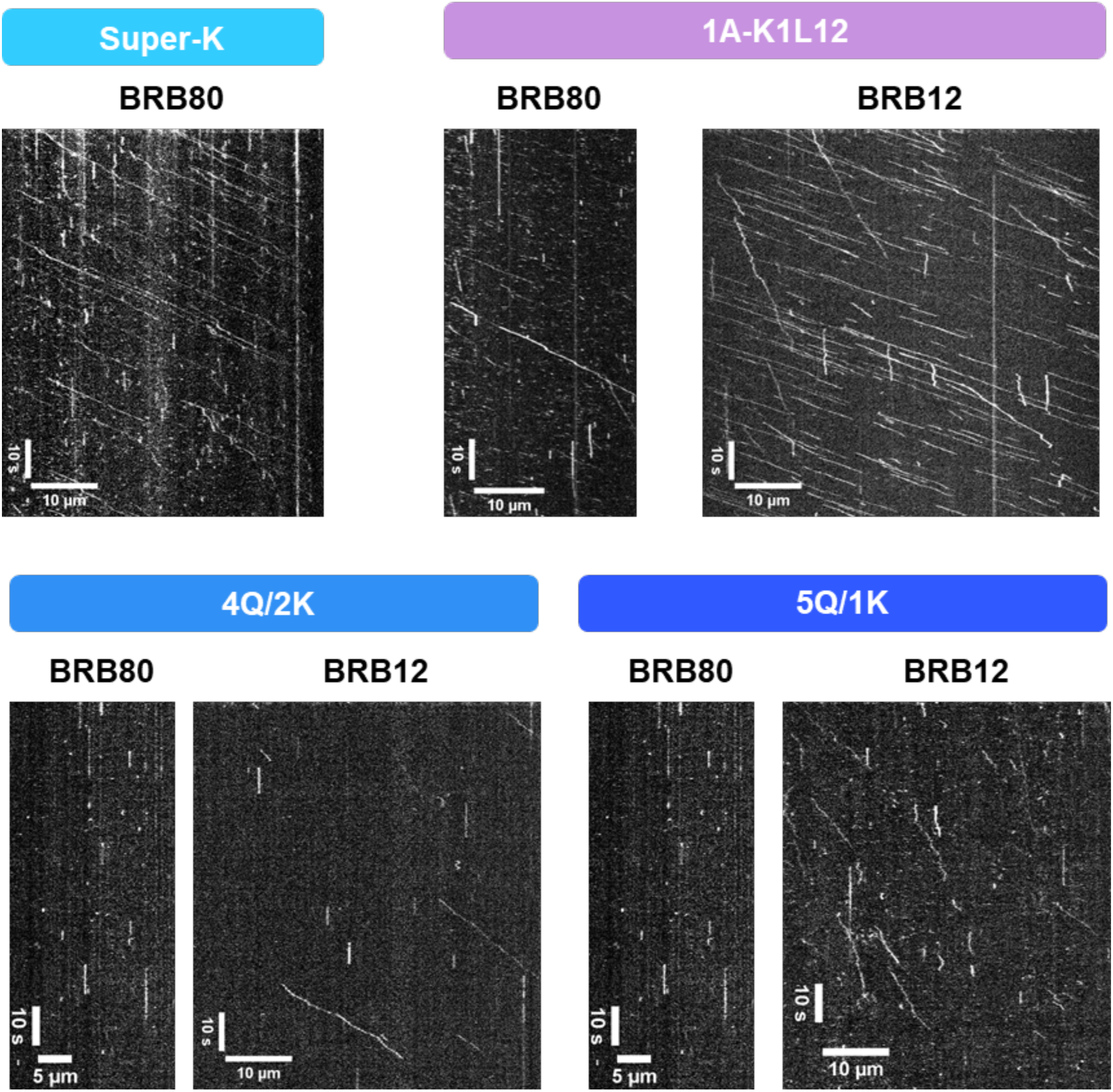
Example Kymographs of Charge Mutants in BRB80 and BRB12. Representative kymographs for the data presented in Figure 4.

**Figure S5:**
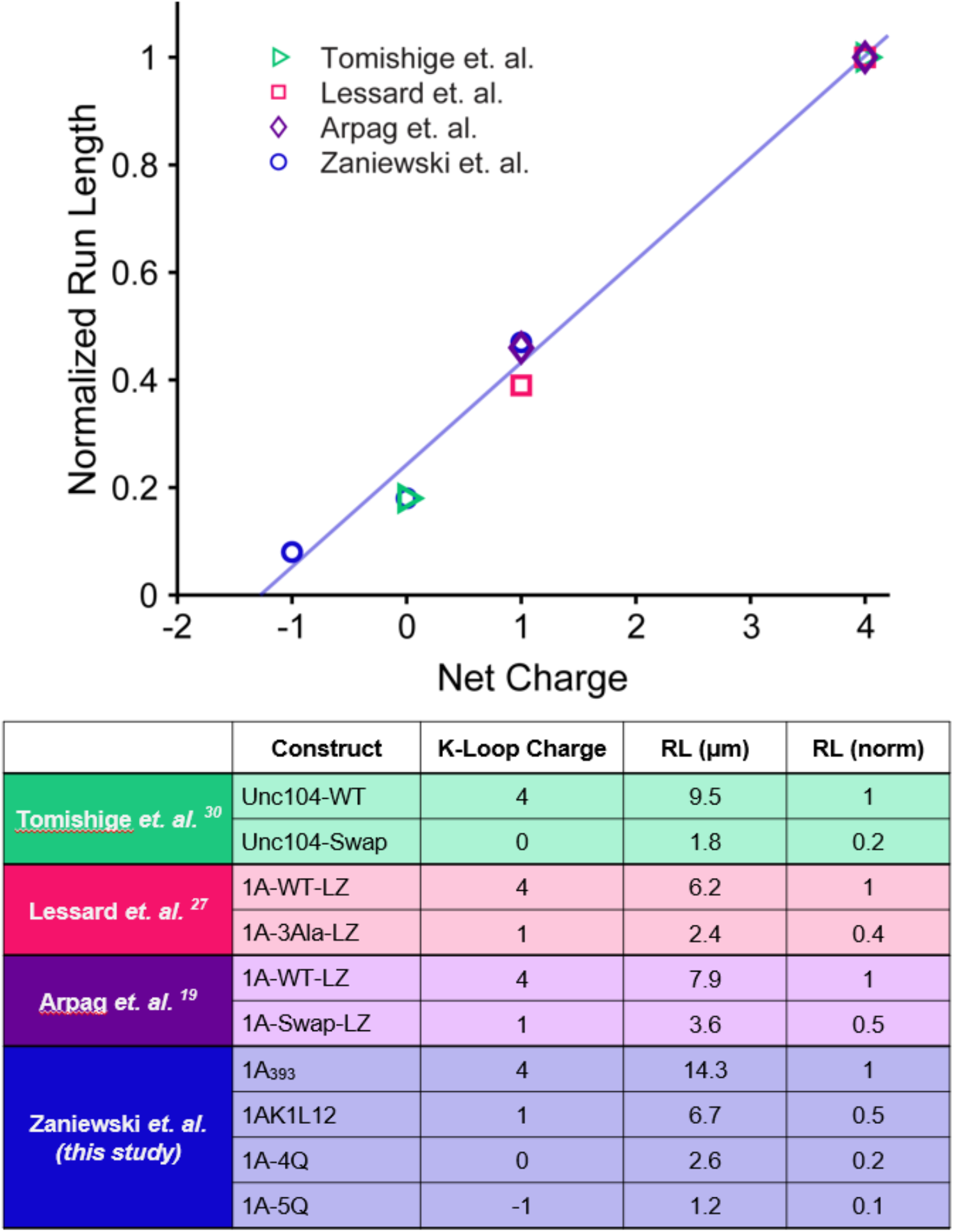
Published run lengths versus K-loop charge for stably dimerized KIF1A constructs in BRB12. (Top) Plot of normalized run length versus net charge of Loop-12 for the present study and three published studies. Run lengths are normalized to the respective wild-type value (+4 net charge) for each study. Line represents fit to the relative run length versus charge for the present study (see Figure 5 for plot of non-normalized data). All experiments were performed in 12 mM PIPES buffer using constitutively active KIF1A dimers stabilized by an added coiled-coil domain. (Bottom) Actual and normalized run lengths from the four studies.

